# Globin digest and its constituent peptides promote skeletal muscle hypertrophy and enhance physical performance

**DOI:** 10.64898/2026.05.27.728339

**Authors:** Misa Nakai, Liqing Zang, Kazutake Fukada, Keiichi Ishido, Norihiro Nishimura, Yasuhito Shimada

**Affiliations:** Graduate School of Regional Innovation Studies, Mie University, 1377 Kurimamachiya-cho, Tsu, Mie 514-8507, Japan; Mie University Zebrafish Research Center, 2-174 Edobashi, Tsu, Mie 514-8507, Japan; Rohto Pharmaceutical Co., Ltd., 1-8-1 Tatsumi-Nishi, Ikuno-ku, Osaka 544-8666, Japan; MG Pharma Inc., 7-7-25 Saito-Asagi, Ibaraki, Osaka 567-0085, Japan; Department of Integrative Pharmacology, Mie University Graduate School of Medicine, 2-174 Edobashi, Tsu, Mie 514-8507, Japan

**Keywords:** Globin digest, bioactive peptides, myogenesis, zebrafish

## Abstract

Globin digest (GD), an acidic protease hydrolysate of hemoglobin, has been recognized for its anti-obesity and glucose-modulating effects; however, its direct anabolic potential in skeletal muscle remains uncharacterized. We evaluated the effects of GD and its constituent peptides on muscle hypertrophy and motor function using zebrafish, mice, and C2C12 myoblasts. Adult zebrafish administered GD (400 mg/kg BW/d) for 1 week showed significantly increased swimming distance (*p* < 0.05). Similarly, oral administration of GD (1 g/kg BW/d) to C57BL/6J mice for 4 weeks enhanced grip strength and rotarod performance, accompanied by a 1.5-fold increase in myofiber diameter and upregulation of fast-twitch *Myh1* (1.9-fold) and *Myh2* (1.8-fold) mRNA levels. *In vitro*, GD dose-dependently (1–100 μg/mL) stimulated C2C12 differentiation and MyHC accumulation. Notably, GD did not merely serve as a nutritional nitrogen source; instead, it functioned as a signaling modulator via a specific “relay-like” peptide orchestration. Among six identified sequences, Peptides 3 (WTQR) and 5 (WGK) primarily initiated early-stage commitment by upregulating *MyoD* and *Myf5*, whereas Peptides 2 (VVYP) and 6 (FES) accelerated mid-stage maturation. This stage-specific synergy achieved robust myotube hypertrophy that exceeded the efficacy of individual components. These findings demonstrate that GD promotes skeletal muscle hypertrophy and motor function through direct myogenic signaling, establishing a novel foundation for precision sports nutrition to optimize muscle maintenance and physical performance.

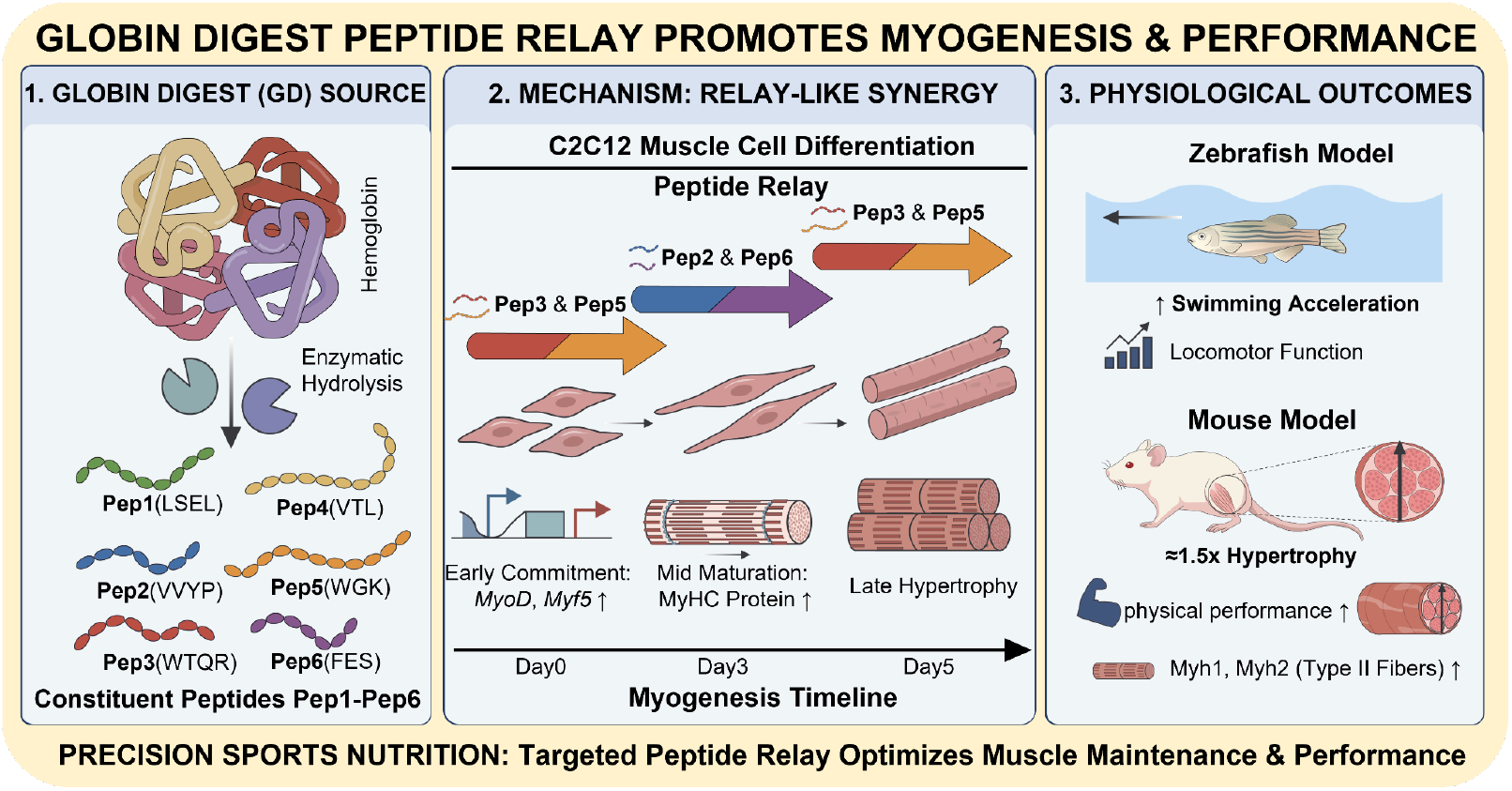

## 1. INTRODUCTION

Optimizing skeletal muscle mass and function is a critical priority for both elite athletes aiming for peak performance and the general population seeking to maintain metabolic health^[1– 3^]. To maximize muscle protein synthesis, current nutritional guidelines recommend a high daily protein intake of 1.6 to 2.2 g/kg body weight for athletes^[4]^. However, achieving these targets through conventional dietary sources while maintaining high bioavailability remains a significant practical challenge. Athletes frequently encounter a “quantity versus quality” trade-off; increasing total protein intake to meet physiological demands can lead to gastrointestinal distress and suboptimal absorption efficiency, ultimately hindering consistent nutritional management^[5–7]^.

To overcome these physiological bottlenecks, nutritional strategies must shift from merely supplying bulk building blocks to efficiently stimulating the skeletal muscle regenerative program. Myogenesis is a highly orchestrated process regulated by myogenic regulatory factors (MRFs). During early differentiation, *Myf5* and *MyoD* proteins drive the commitment and proliferation of myogenic progenitor cells, followed by the upregulation of muscle-specific myosin heavy chain (*MyHC*) genes (*Myh1, Myh2*), which leads to the functional maturation and hypertrophy of fast-twitch (Type II) fibers ^[8–10]^. Because Type II fibers are critically responsible for explosive power and athletic performance^[11,12]^, compounds that directly trigger this Myf5- or MyoD-mediated pathway are highly sought after. In this context, bioactive short-chain peptides have emerged as a potent alternative to intact proteins^[13–16]^. They are rapidly internalized into cells via independent, high-capacity peptide transporters such as PepT1^[17]^ and PHT1^[18]^ and function as powerful signaling ligands capable of inducing MRF expression at significantly lower doses^[19,20]^.

Food-derived bioactive oligopeptides have gained considerable attention for their diverse biological activities, including anti-inflammatory, anti-fatigue, and metabolic regulatory effects^[21–23]^. Among these, globin digest (GD), obtained through the acidic protease treatment of porcine hemoglobin, is approved as a functional food in Japan and China^[24]^. GD provides potent metabolic benefits, including improvements in postprandial hyperlipidemia and hyperglycemia^[25,26]^. Our recent studies demonstrated that GD exerts anti-obesity effects by upregulating uncoupling protein 1 (UCP1)-related pathways and enhancing systemic energy expenditure. Furthermore, we previously demonstrated that GD upregulates the expression of *Ucp2* and *Ucp3* genes specifically within skeletal muscle, where they are thought to enhance fatty acid oxidation and energy expenditure. Crucially, we observed that GD-treated *in vivo* models exhibited a marked reduction in visceral fat without a corresponding decrease in total body weight, a phenomenon accompanied by preliminary evidence of upregulated myogenic biomarkers^[27]^. These findings led to the hypothesis that the anti-obesity effects of GD are complemented by a concomitant increase in muscle mass, pointing to a beneficial modulation of body composition through the adipo-muscular axis. Despite these promising systemic observations, the direct cellular mechanisms by which GD modulates the myogenic program and determines muscle fiber-type composition remain uncharacterized. Furthermore, the specific peptide sequences within the GD mixture responsible for these direct anabolic effects have not been identified. Therefore, determining whether GD optimizes the intramuscular microenvironment to support Type II fiber hypertrophy represents a novel paradigm in sports nutrition.

In the present study, we employed an integrated *in vivo* and *in vitro* approach to evaluate the effects of GD on skeletal muscle differentiation and motor function. First, we assessed motor function improvements in zebrafish, followed by an evaluation of exercise performance and the expression of key myogenic genes (*Myf5, MyoD, Myh1, Myh2, Myh3, and Myh7*) in a mouse model. Furthermore, to elucidate the underlying molecular mechanisms, we investigated the direct effects of GD on myogenesis and MyHC assembly using C2C12 myoblasts. Additionally, we identified six specific peptide sequences within GD and characterized their individual roles in promoting myogenic differentiation. These findings establish the multifaceted nutritional value of GD and provide a foundation for novel nutritional strategies to support efficient muscle maintenance and performance enhancement in athletes.

## 2. MATERIALS AND METHODS

### 2.1. Ethics statement

All animal procedures were approved by the Ethics Committee of Mie University, Tsu, Japan (Approval No. 2024-19). Animal experiments were performed in accordance with the Japanese Animal Welfare Regulatory Practice Act on Welfare and Management of Animals (Ministry of the Environment, Japan) and complied with international guidelines.

### 2.2. GD and its constituent peptides

This study utilized globin digest (GD; MG Pharma, Osaka, Japan), an enzymatic hydrolysate of porcine hemoglobin, was used in this study^[28]^. The six constituent peptides of GD evaluated in this study—LSEL (Leucine-Serine-Glutamic acid-Leucine), VVYP (Valine-Valine-Tyrosine-Proline), WTQR (Tryptophan-Threonine-Glutamine-Arginine), VTL (Valine-Threonine-Leucine), WGK (Tryptophan-Glycine-Lysine), and FES (Phenylalanine-Glutamic acid-Serine) —were synthesized by GenScript (Tokyo, Japan). The origins of these peptide sequences within porcine hemoglobin and their respective quantitative contents in GD are summarized in **Table 1**.

**Table 1.** The sequences, origins, and contents of the six constituent peptides within GD.

| Peptide | Peptide sequence | Origin in porcine hemoglobin | Reference |
| --- | --- | --- | --- |
| Pep 1 | LSEL | β-chain (89–92) | [25] |
| Pep 2 | VVYP | β-chain (33–36) | [24] |
| Pep 3 | WTQR | β-chain (37–40) | [28] |
| Pep 4 | VTL | α-chain (107–109) | [24] |
| Pep 5 | WGK | α-chain (14–16) and β-chain (15–17) | [29] |
| Pep 6 | FES | β-chain (42–44) | [29] |

### 2.3. Cell culture

The C2C12 mouse myoblast cell line (RCB0987) was purchased from the RIKEN BRC through the National Bio-Resource Project of the MEXT, Japan. The cells were cultured in a 37 °C, 5% CO_2_ incubator using a growth medium composed of Dulbecco’s Modified Eagle’s Medium (DMEM; Sigma-Aldrich, St. Louis, MO, USA) supplemented with 10% Fetal Bovine Serum (FBS; Sigma-Aldrich) and 1% Antibiotic-Antimycotic (Nacalai Tesque, Kyoto, Japan). When the cells reached 70% confluency, the culture medium was replaced with differentiation medium (DMEM supplemented with 2% horse serum [HS; Gibco, Grand Island, NY, USA]). The differentiation medium was refreshed every other day. GD (1, 10 or 100 μg/mL) and individual peptides (1 μg/mL) were added at the onset of differentiation.

### 2.4. Animals and husbandry

Zebrafish (*Danio rerio*, AB line) were purchased from the Zebrafish International Research Center (ZIRC, Eugene, OR, USA) and maintained at our facility under standard laboratory conditions. The zebrafish were fed Gemma Micro 75, 150, and 300 (Skretting, Fontaine-les-Vervins, France) according to their developmental stages or body length. Wild-type C57BL/6J mice were purchased from Japan SLC (Hamamatsu, Japan) and housed at the Institute of Laboratory Animals at Mie University.

### 2.5. Zebrafish experiment

For the oral administration of GD to adult zebrafish, zebrafish food containing 10% (w/w) GD was prepared using gluten as a carrier material, as previously described^[30]^. Briefly, 3-month-old female zebrafish were randomly assigned to either the control or GD group (five fish per 2-L tank). The control group was fed a normal diet (gluten granules; 2 mg/fish/day) throughout the 1-week experimental period. The GD group was fed GD-containing gluten granules (2 mg/fish/day), corresponding to a dose of 400 mg/kg body weight/day. At the end of the feeding experiment, body weights were measured, followed by the electrically stimulated swimming test. For this test, individual zebrafish were placed in a rectangular tank (17.5 cm × 14.5cm) and allowed to acclimate for at least 5 min. Once the fish ceased spontaneous movement and became completely motionless, a 5-V electrical pulse was applied for 0.1 s from electrodes positioned at both ends of the tank to induce swimming. The experimental arena was filmed from directly above at 30 frames per second using a digital high-definition (HD) video camera (HC-V495M; Panasonic, Osaka, Japan). The stimulus-evoked swimming speed, acceleration, and distance traveled were subsequently analyzed using ToxTrac software (version 1.1)^[31]^.

### 2.6. Mouse experiment

Six-week-old male C57BL/6JmsSlc mice were fed a diet containing 0.5% GD for 4 weeks under ad libitum conditions. Based on the average daily food intake (approximately 4 g/day) and average body weight (approximately 20–23 g), the estimated daily dose of GD was 1 g/kg body weight/day. Six to seven mice were housed per cage. Body weight, all-limb grip strength, rotarod performance, and residual feed weight were recorded weekly. Grip strength was measured using a horizontal-type small animal grip strength meter (GPM-101B; Melquest, Toyama, Japan), allowing for the simultaneous measurement of all four limbs. Motor coordination and endurance were evaluated using a mouse rotarod apparatus (RTR-M1; Melquest). The test was performed using an accelerating protocol from 4 to 40 rpm over 300 s (The Jackson Laboratory Mouse Neurobehavioral Phenotyping Facility protocol)^[32]^. From the second week onward, mice that fell in the same direction as the rotation (abnormal adaptive movement) were excluded from the analysis. At the end of the experiment, mice were euthanized by over-anesthesia with isoflurane (Pfizer, Pearl River, NY, USA). Gastrocnemius muscle tissues were dissected for subsequent histology and RT-qPCR analysis.

### 2.7. Histological analysis

Dissected tissues were fixed overnight in 4% paraformaldehyde in phosphate-buffered saline (4% PFA; Falma, Tokyo, Japan) at 4 °C. The fixed samples were sent to the Division of Medical Research Support, Advanced Research Support Center (ADRES), Ehime University, where they were processed for paraffin embedding, sectioning (5-μm thickness), and routine hematoxylin and eosin (H&E) staining. Stained sections were imaged in brightfield mode using a BZ-X710 fluorescence microscope (Keyence, Osaka, Japan). To quantify muscle fiber hypertrophy, morphometric analyses were performed using ImageJ software (Fiji distribution, version 1.52p; National Institutes of Health, Bethesda, MD, USA). For the zebrafish trunk muscle, sections displaying the optimal transverse orientation of the muscle were selected, and the transverse diameter of the myofibers was measured at five distinct locations per animal, with the average representing the individual value. For the mouse gastrocnemius muscle, the cross-sectional area of at least 20 individual myofibers per animal was directly measured from the H&E-stained transverse sections to evaluate muscle hypertrophy.

### 2.8. Quantitative reverse transcription PCR (qRT-PCR)

Mouse tissues and C2C12 cells were homogenized in TRIzol reagent (Invitrogen, Carlsbad, CA, USA) using a mechanical homogenizer. The homogenized samples were subjected to phase separation with chloroform, and the aqueous phase was transferred to RNeasy columns for further purification using the RNeasy Mini Kit (74104; QIAGEN, Hilden, Germany) according to the manufacturer’s instructions. The purity and the concentration of the extracted RNA were determined by measuring the absorbance ratio at 260 nm and 280 nm. cDNA was synthesized from 400 ng of total RNA using the ReverTra Ace qPCR RT Master Mix with gDNA Remover (TOYOBO, Osaka, Japan). Quantitative RT-PCR was performed using Power SYBR Green Master Mix (Applied Biosystems, Foster City, CA, USA) and the ABI StepOnePlus Real-Time PCR System (Applied Biosystems) in accordance with the manufacturer’s instructions. Relative mRNA expression levels were quantified and normalized to *Hprt* as an endogenous control gene. The detailed primer sequences used for amplification are listed in **Supplementary Table S1**.

### 2.9. Immunofluorescence analysis

C2C12 myotubes were washed three times and fixed with 4% paraformaldehyde for 10 min at 4°C. After fixation, the cells were washed three times with ice-cold PBS and permeabilized using 0.1% Triton X-100 (Sigma-Aldrich) for 10 min. To minimize non-specific binding, the cells were blocked with PBST (PBS containing 0.1% Tween 20) supplemented with 1% BSA (Sigma-Aldrich) and 22.52 mg/mL glycine for 2 h. The cells were then incubated with a primary antibody against myosin heavy chain (MAB4470, 1:200 dilution; R&D Systems, Minneapolis, MN, USA) overnight at 4°C. The solution was decanted, and the cells were washed three times in PBS for 5 min each. Myotubes were then treated with a secondary antibody (Anti-mouse IgG (H+L), F(ab’)2 Fragment, Alexa Fluor® 594 Conjugate; #8890S; Cell Signaling Technology, Danvers, MA, USA; 1:1000 dilution) in 1% BSA in PBST for 1 h at room temperature in the dark. Nuclei were stained using Hoechst 33342 (1:1000 dilution; Molecular Probes, Eugene, OR, USA) for 10 min. After washing with PBS, images of the stained C2C12 myotubes were captured using a fluorescence microscope (BZ-X710; Keyence, Osaka, Japan). Image analysis and quantification were performed using ImageJ software.

### 2.10. Statistical Analysis

All data were analyzed using Student’s *t*-test, Welch’s *t*-test, or one-way analysis of variance (ANOVA) followed by the Bonferroni–Dunn multiple comparison procedure, depending on the number of comparisons. Analyses were performed using GraphPad Prism version 11 (GraphPad Software, San Diego, CA, USA). Results with *p* < 0.05 were considered statistically significant.

## 3. RESULTS

### 3.1. GD promotes muscle hypertrophy and locomotor ability in zebrafish

To evaluate the *in vivo* effects of GD on skeletal muscle performance, we first utilized a zebrafish model. Young zebrafish (3 mpf) were orally administered GD (400 mg/kg body weight) for one week. Prior to administration, the standard length (SL) and body weight (BW) of the control group were 2.0 ± 0.1 cm and 0.11 ± 0.02 g, respectively, while those of the GD group were 1.9 ± 0.1 cm and 0.09 ± 0.01 g. After the 1-week feeding trial, the SL increased to 2.1 ± 0.1 cm in the control group and 2.0 ± 0.2 cm in the GD group, representing a 5% increase in both groups. Concurrently, the BW increased to 0.13 ± 0.02 g in the control group (a 13% increase) and 0.12 ± 0.01 g in the GD group (a 19% increase). Although the GD-treated group exhibited a greater percentage of weight gain, no statistically significant differences were observed in either SL or BW between the groups (**Figure S1**). Following the treatment, an electrically stimulated forced swimming test was conducted to assess functional output. As shown in **Figure 1**, the GD-treated group demonstrated robust improvements across multiple kinematic parameters. Specifically, the average acceleration during the escape response was significantly (*p* < 0.05) increased by 2.9-fold compared with the control group (**Figure 1A**). Similarly, the maximum burst acceleration was markedly elevated by 3.8-fold (**Figure 1B**, *p* < 0.05). This pronounced enhancement indicates that GD administration specifically amplifies the explosive motor power of zebrafish. Consistent with the elevated acceleration, the average swimming speed was also significantly higher, showing a 90% increase in the GD-treated group (**Figure 1C**, *p* < 0.05). To investigate whether this functional enhancement was associated with structural changes in the musculature, we performed a histological analysis using hematoxylin and eosin (HE) staining of the trunk skeletal muscle **(Figure 1D)**. Representative images revealed that zebrafish treated with GD exhibited visibly larger skeletal muscle fibers than those in the control group. Morphometric analysis showed that the muscle fiber diameter in the GD-treated group was 1.39-fold thicker than that of the control group (**Figure 1E**, *p* < 0.01).

**Figure 1.**
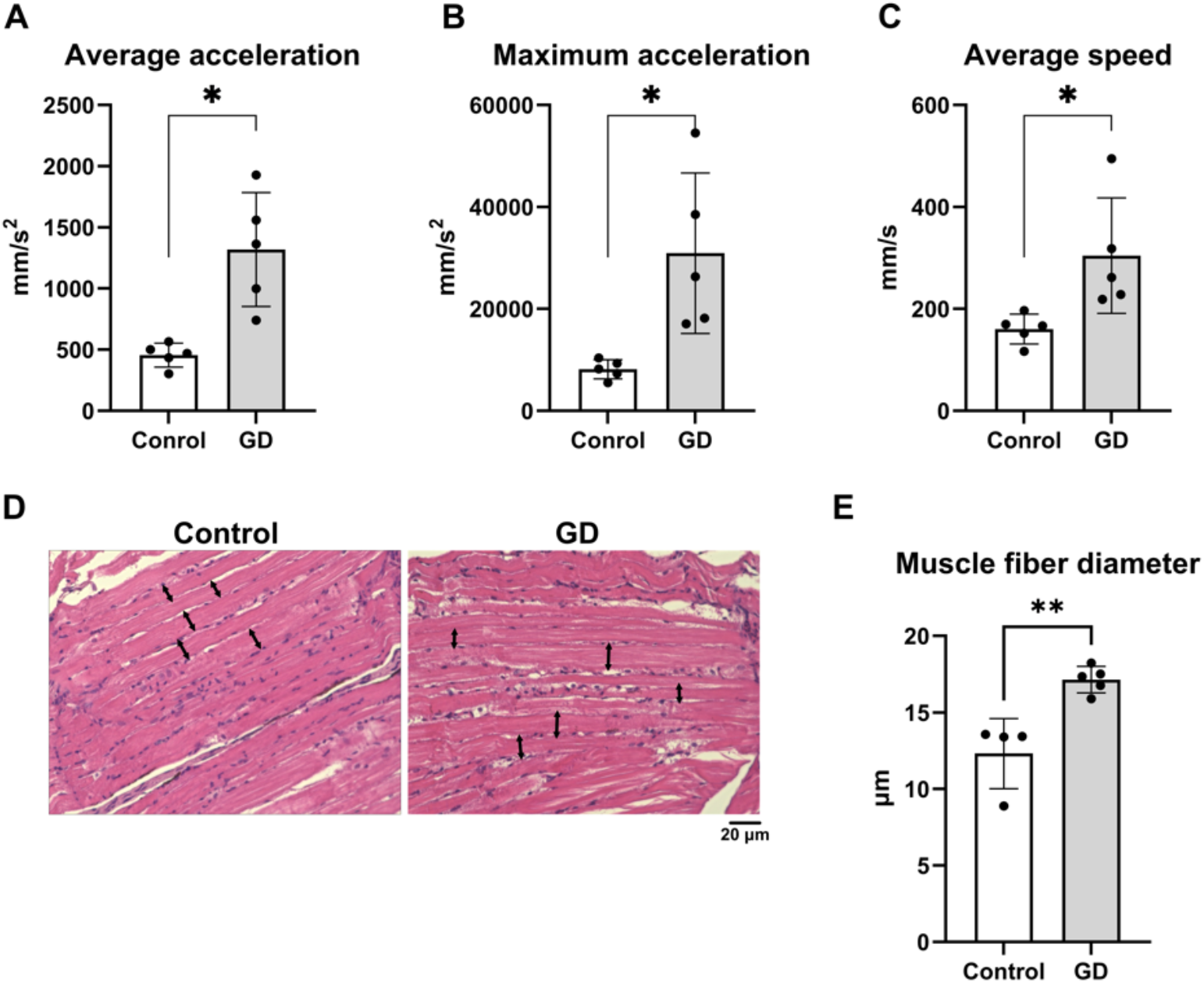
Effects of GD administration on burst locomotor performance and muscle fiber morphology in young adult zebrafish. Kinematic analysis of the electrically stimulated forced swimming test showing (**A**) average acceleration, (**B**) maximum acceleration, and (**C**) average speed. All parameters were significantly enhanced in the GD-treated group. Data are presented as mean ± standard deviation (SD). \**p* < 0.05, n = 5. (**D**) Representative histological images of H&E stained sections of the trunk skeletal muscle. Black arrows indicate the transverse diameter of individual muscle fibers. (**E**) Quantitative morphometric analysis of the skeletal muscle fiber diameter. For each animal, the transverse diameter was measured at five distinct locations from the optimal cross-section, and the average was used as the individual value. Data are presented as mean ± SD. \*\**p* < 0.01, (control: n = 4, GD: n = 5).

### 3.2. GD enhances skeletal muscle strength and promotes myofiber hypertrophy in mice

To examine whether the skeletal muscle enhancing effects of GD observed in zebrafish are conserved in mammals, we orally administered GD to mice for four weeks and conducted a series of physical function tests. During the administration period, both the control and GD-treated groups exhibited normal growth. Prior to administration, the body weights of the control and GD groups were 20.7 ± 0.6 g and 20.7 ± 0.5 g, respectively. After four weeks, the weights increased to 23.1 ± 0.8 g in the control group and 21.8 ± 1.9 g in the GD group, with no significant differences observed between the groups (**Figure S2**). Similarly, the average daily food intake per mouse was comparable (4.13 g for control vs. 4.06 g for GD; **Figure S3**).

Despite this comparable body mass and energy intake, and consistent with the findings in zebrafish, GD administration significantly improved muscle strength and coordination. Specifically, the GD group demonstrated a clear and statistically significant superiority in both grip strength (an 14.4% increase) and rotarod latency (an 1.79-fold increase) by week 4 (**Figure 2A** and **2B**, *p* < 0.05). Notably, at the 4-week mark, several mice in the GD-treated group demonstrated exceptional grip and balance, remaining attached to the rotating rod and maintaining their position even during full 360° rotations, a level of performance not observed in the control group. To determine the structural basis for these functional improvements, we performed a histological analysis of the gastrocnemius muscle using H&E staining **(Figure 2C)**. Morphological examination revealed prominent enlargement of the skeletal muscle fibers in the GD-treated group. Quantitative analysis confirmed these observations, showing that GD administration led to an 1.38-fold increase in myofiber diameter compared with the control group **(Figure 2D)**. Furthermore, to elucidate the molecular changes underlying this hypertrophy, we examined the mRNA expression of myosin heavy chain isoforms *Myh1* and *Myh2* in the gastrocnemius muscle **(Figure 3A and 3B)**. GD administration significantly upregulated the expression of both isoforms compared with the control group (*p* < 0.05). Specifically, the mRNA levels of *Myh1* and *Myh2* increased by 1.90-fold and 1.77-fold, respectively. Furthermore, other myosin isoforms, including *Myh3* and *Myh7*, also showed a strong upward trend in their expression levels (*p* = 0.054 and *p* = 0.053, respectively; **Figure 3C and 3D**). While these changes did not reach conventional statistical significance, the overall elevation across multiple myosin heavy chain genes further supports the robust myogenic effect of GD administration. These molecular data, highlighting a broad increase in contractile protein isoforms (particularly the fast-twitch-associated *Myh1* and *Myh2*), are consistent with the enhanced strength and fiber enlargement observed in GD-treated mice. Collectively, these results demonstrate that GD effectively promotes muscle hypertrophy and enhances physical performance in a mammalian model.

**Figure 2.**
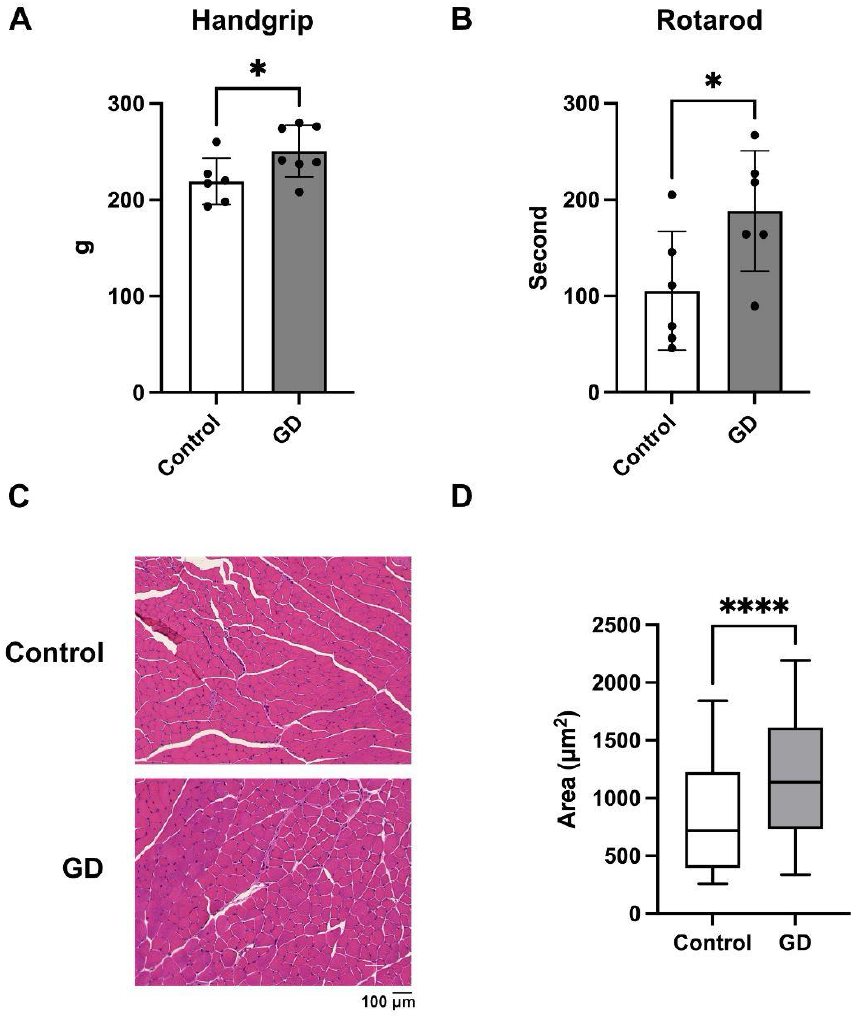
Effects of GD administration on physical performance, muscle morphology, and myogenic gene expression in mice. (**A**) Grip strength of mice during the 4-week administration period (control vs. GD, 4 weeks: *p* < 0.05). **(B)** Latency to fall in the accelerating rotarod test (*p* < 0.05). **(C)** Representative histological images of H&E stained sections of the gastrocnemius muscle. **(D)** Quantitative morphometric analysis of the muscle fiber cross-sectional area (CSA). The CSA of at least 20 individual myofibers per animal was measured. Data are presented as mean ± SD. \**p* < 0.05, \*\*\*\**p* < 0.0001 vs. control. Control: n = 6; GD: n = 7.

**Figure 3.**
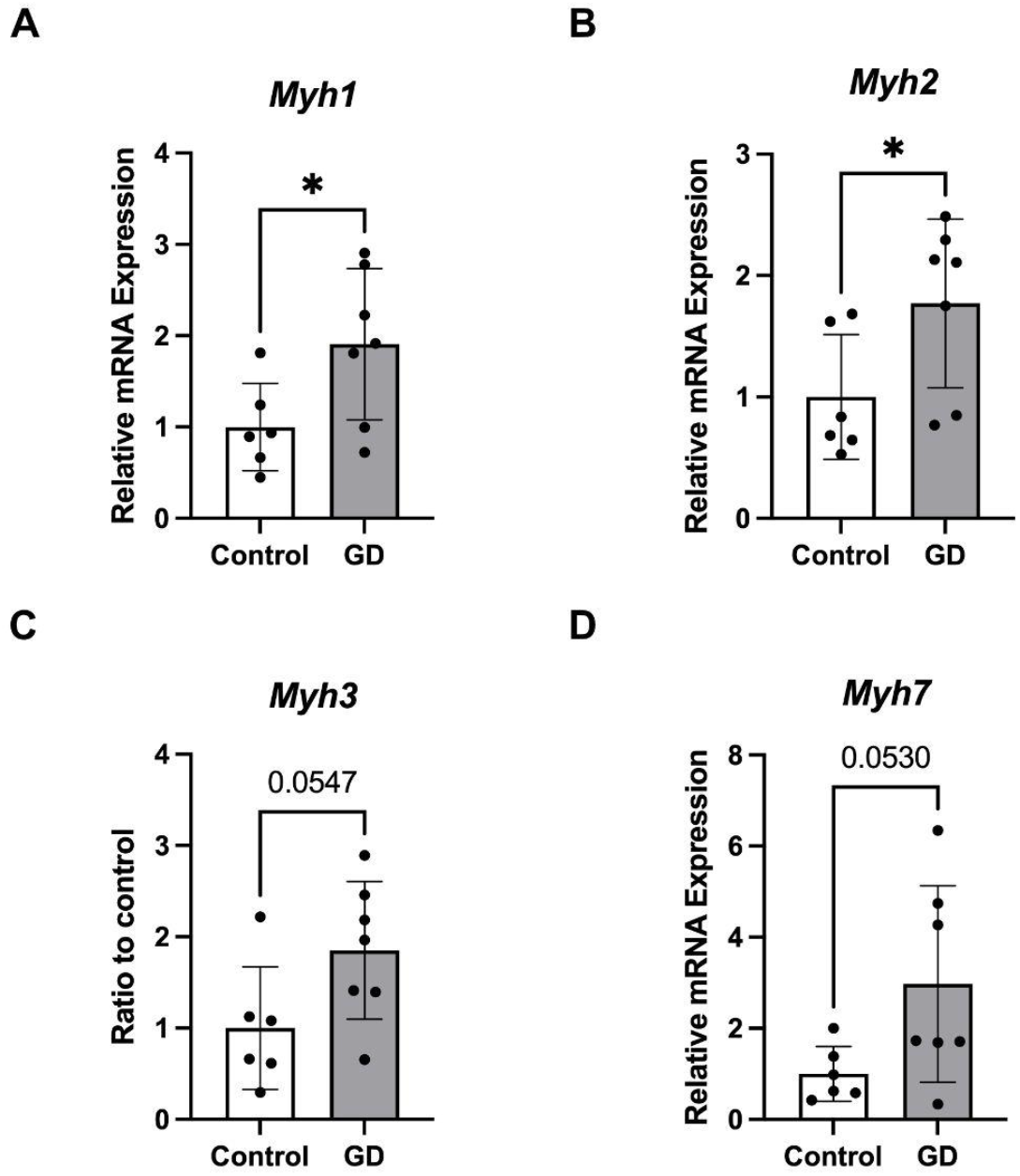
Effects of GD administration on myogenic gene expression in vivo. Relative mRNA expression levels of myosin heavy chain isoforms: (**A**) *Myh1* (*p* < 0.05), (**B**) *Myh2* (*p* < 0.05), (**C**) *Myh3* (*p* = 0.054), and (**D**) *Myh7* (*p* = 0.053). Data are presented as mean ± SD. \**p* < 0.05 vs. control. Control: n = 6; GD: n = 7.

### 3.3. GD and its constituent peptides promote myogenic differentiation in C2C12 cell

Skeletal muscle regeneration and hypertrophy rely on the sequential proliferation and differentiation of myoblasts. We first examined the effect of GD on myoblast proliferation. Treatment of undifferentiated C2C12 cells with GD resulted in a significant, dose-dependent increase in cell proliferation at concentrations of 10 μg/mL and above (**Figure S4A**). To identify the specific bioactive components responsible for the myogenic effects of GD, we focused on six short-chain peptides (Peptide 1–6: LSEL, VVYP, WTQR, VTL, WGK, and FES; **Table 1**) previously identified as key constituents of GD^[24,25,28,29]^. Among these peptides tested individually at 1 μg/mL, Peptides 2 and 3 significantly stimulated cell proliferation compared with the control (**Figure S4B**).

Following this initial expansion phase, we investigated the direct cellular effects of GD on myogenic differentiation *in vitro*. C2C12 myoblasts were cultured in differentiation medium with or without GD for day 5. Immunofluorescence analysis revealed a robust increase in the expression of myosin heavy chain (MyHC), a late-stage marker of myogenic maturation, in GD-treated cells compared with control cells (**Figure 4A**). The treatment with GD markedly enhanced the morphological maturation of the cells, as evidenced by the increased formation of MyHC-positive multinucleated myotubes. Quantitative analysis of the immunofluorescence images demonstrated a highly significant increase in the MyHC-positive area in the GD-treated group compared with the control group (*p* < 0.0001). Furthermore, we examined the dose-response relationship of this effect. GD treatment increased MyHC fluorescence intensity in a concentration-dependent manner at 1, 10, and 100 μg/mL **(Figure 4B)**. C2C12 myoblasts were then treated with either GD or each of the six key peptides to compare their individual effects on myotube maturation. To evaluate the temporal effects of these six peptides on myogenic differentiation, MyHC immunofluorescence was performed at two different time points: day 3 and day 5 (**Figure 4C-F**). Quantitative analysis of the MyHC-positive area (Figure 4D) revealed that at day 3, Peptides 2 and 6 significantly promoted myotube formation compared with the control. By day 5, the myogenic effect became more pronounced; all peptides except Peptide 4 showed higher MyHC expression levels than the control, with Peptides 3 and 5 exhibiting particularly robust and significant increases (Figure 4F). To further elucidate the molecular mechanisms underlying these observations, we analyzed the expression of genes involved in skeletal muscle differentiation (**Figure 5**). Consistent with the morphological data, the mRNA levels of early myogenic regulatory factors, *Myf5* (**Figure 5A**) and *MyoD* (**Figure 5B**), were upregulated by most peptides (except Peptide 4) compared with the control. Notably, Peptides 3 and 5 induced the most substantial increase in *MyoD* expression. Interestingly, while MyHC protein accumulation was significantly elevated at day 5, the mRNA levels of late-stage markers such as *Myog* (**Figure 5C**), *Myh1*(**Figure 5D**) *and Myh2* (**Figure 5E**) showed no significant differences at this same time point. This discrepancy likely reflects the temporal gap between transcription and translation, the peak of mRNA expression for these markers presumably occurred earlier, leading to the robust protein levels observed during the terminal stage of differentiation.These results suggest that these bioactive peptides, particularly Peptides 3 and 5, may primarily accelerate the early stages of the myogenic program by modulating *MyoD* and *Myf5* expression, leading to the enhanced MyHC protein accumulation observed in later stages.

**Figure 4.**
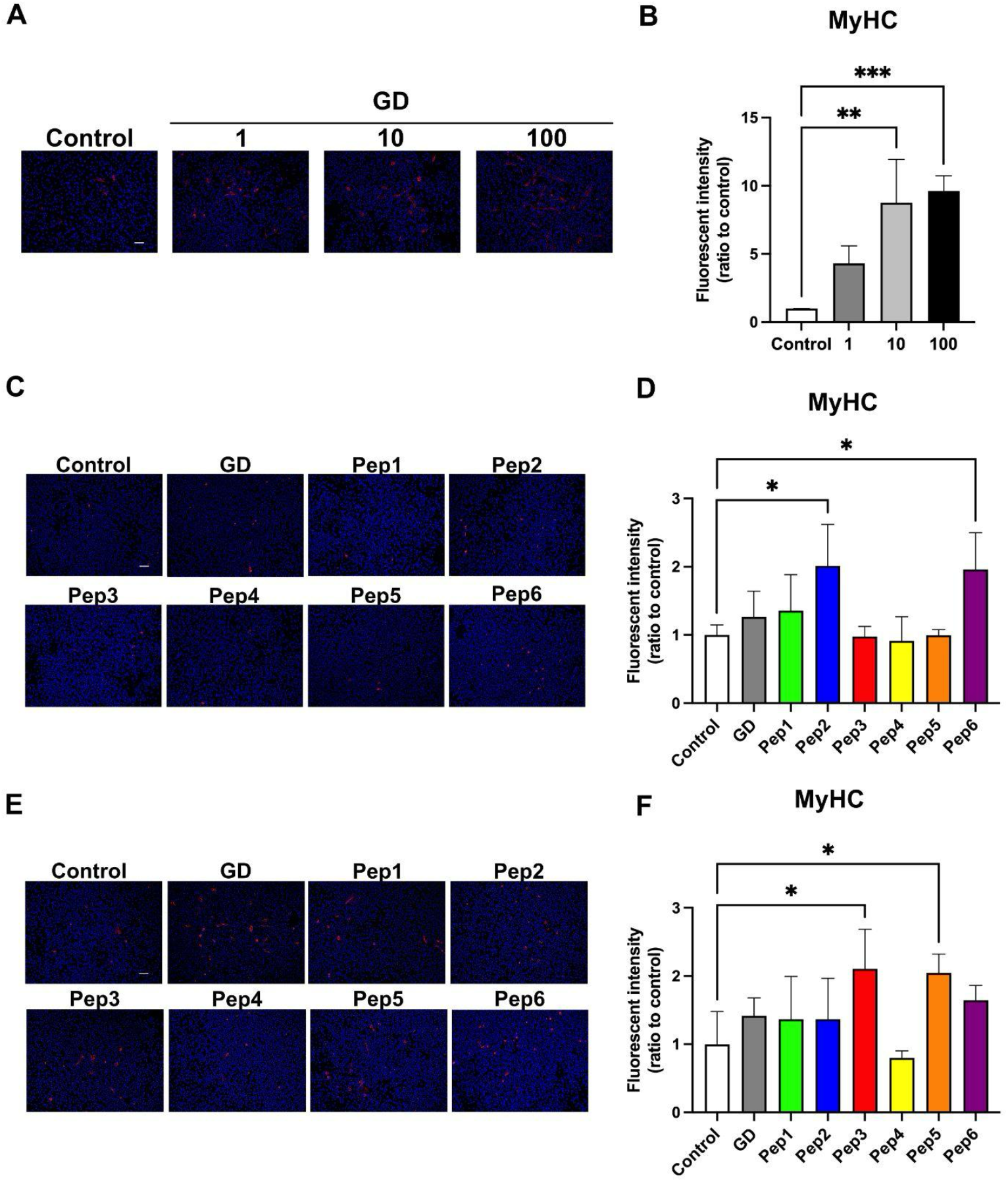
Effects of GD and its constituent bioactive peptides on myogenic differentiation in C2C12 myoblasts. **(A)** Representative immunofluorescence images of myosin heavy chain (MyHC) expression in C2C12 myotubes after 5 days of differentiation with or without GD. **(B)** Dose-response analysis of MyHC fluorescence intensity in C2C12 cells treated with GD (1, 10, and 100 μg/mL). **(C)** Representative MyHC immunofluorescence images and **(D)** quantitative analysis of MyHC-positive area at day 3 of differentiation following treatment with Peptides 1–6 (Control vs. Pep 2 or 6: *p* < 0.05). **(E)** Representative MyHC immunofluorescence images and **(F)** quantitative analysis of MyHC-positive area at day 5 (Control vs. Pep 3 or 5: *p* < 0.05). Data are presented as mean ± SD. \**p* < 0.05, \*\*\*\**p* < 0.0001 vs. control, n = 5. Scale bar = 100 μm.

**Figure 5.**
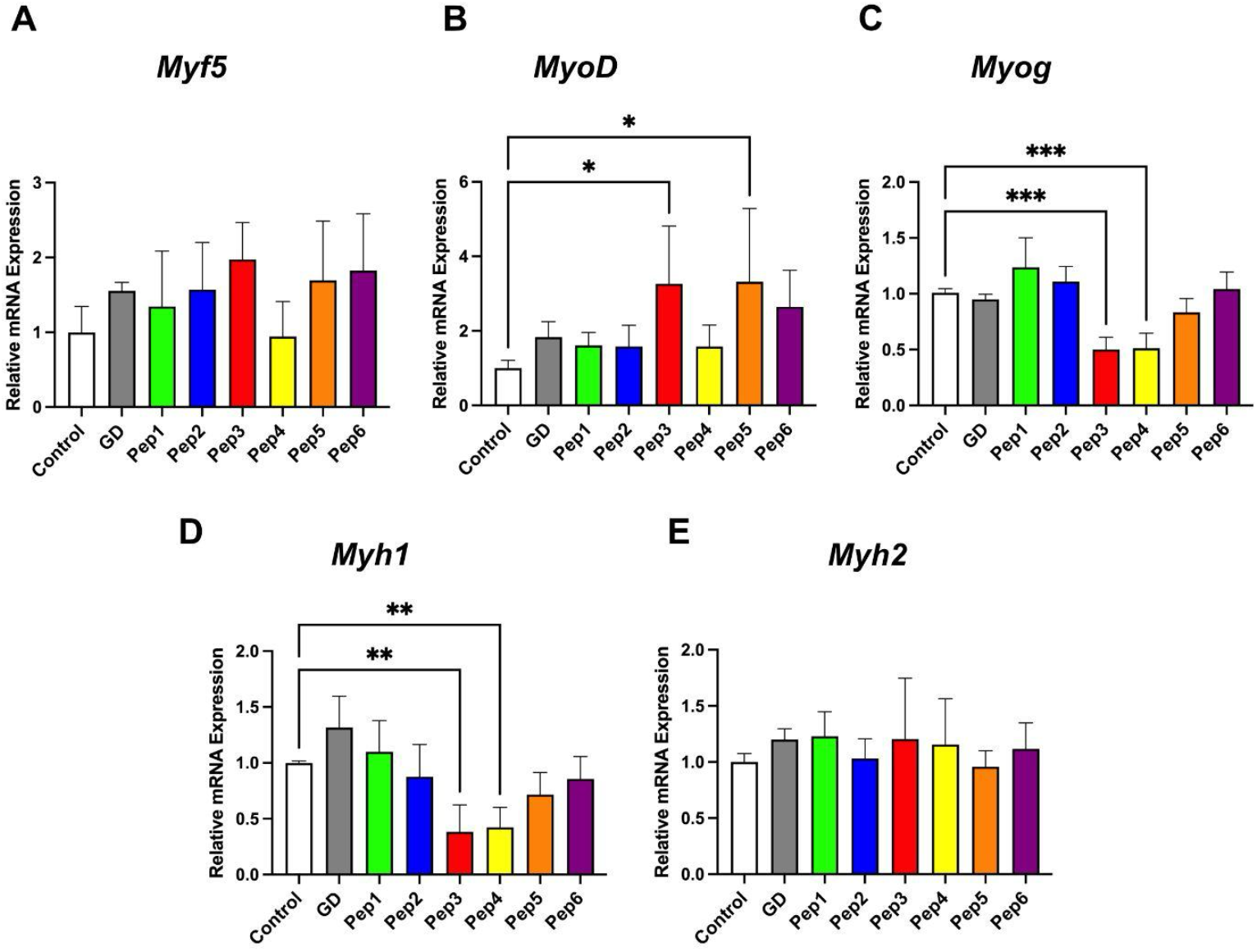
GD-derived peptides modulate the expression of myogenic regulatory factors and maturation markers. Relative mRNA expression levels of myogenic regulatory factors, **(A)** *Myf5*, **(B)** *MyoD*, **(C)** *Myog* at day 3 and maturation markers **(D)** *Myh1*, **(E)** *Myh2* at day 5 in C2C12 cells. Data are presented as mean ± SD. \**p* < 0.05, \*\**p* < 0.01, \*\*\*\**p* < 0.0001 vs. control, n = 5.

## 4. DISCUSSION

In the present study, we demonstrated that globin digest (GD) and its specific constituent peptides directly promote skeletal muscle hypertrophy and enhance physical performance. Our results revealed that GD administration significantly increases muscle fiber diameter and enhances explosive motor function across different species. Furthermore, the identification of a stage-specific peptide relay provides a novel mechanistic explanation for the potent anabolic effects of the natural GD mixture.

### 4.1. Functional expansion of GD: The adipo-muscular axis

Our previous research established that GD exerts anti-obesity effects by upregulating UCP1-mediated thermogenic pathways and suppressing adipocyte hypertrophy in zebrafish and mouse models^[27]^. Our previous findings highlighted that the metabolic impact of GD is mediated not only by *UCP1* in adipose tissue but also by the systemic activation of other UCP family members, including *Ucp2* and *Ucp3* in skeletal muscle. Since *UCP3* is primarily expressed in skeletal muscle and is known to protect mitochondria from oxidative stress during fatty acid oxidation while increasing energy expenditure within myocytes^[39–41]^, the potentiation of myogenic differentiation observed in this study may synergistically enhance these UCP-mediated metabolic benefits.

Given the fundamental role of the adipose-muscle crosstalk in systemic metabolic homeostasis^[42–44]^, these findings provide a molecular basis for the hypothesis that GD acts as a multi-target nutrient. Notably, GD-treated obese models in our previous study exhibited a significant reduction in visceral adipose tissue without a corresponding loss in total body weight^[27]^. At that time, we hypothesized that this maintenance of body mass was due to a concomitant increase in muscle volume. Crucially, the present study confirmed that GD administration induces hypertrophy of skeletal muscle fibers (Figure 2D) and upregulates the expression of myogenic markers, including *Myh1* (Figure 3A) and *Myh2* (Figure 3B) in vivo. By demonstrating that GD directly acts on muscle progenitors to facilitate their maturation, the current C2C12 results provide the crucial mechanistic link (Figure 5), confirming that the observed muscle preservation is driven by the direct enhancement of myogenesis.

The metabolic benefits of GD are not confined to adipose tissue but extend to the functional improvement of skeletal muscle, potentially reflecting the integrated metabolic crosstalk between these tissues. Furthermore, GD administration was previously shown to upregulate *Prkaa2* (AMPK) and *Bdnf*^*[27]*^, both of which are critical regulators of lipid catabolism and the maintenance of muscle fiber integrity^[45,46]^. This dual functionality aligns with recent trends in molecular nutrition, where bioactive molecules such as GLP-1 receptor agonists^[47]^ and Angiotensin-(1-7)^[48]^ are identified as significant modulators of both lipid metabolism and muscle regenerative capacity. The ability of GD to facilitate myogenesis while simultaneously ameliorating visceral adiposity suggests a promising nutritional strategy targeting the adipo-muscular axis, providing a comprehensive intervention for both the clinical management of sarcopenic obesity^[49]^ and the enhancement of athletic power-to-weight ratios to maintain physical independence across the lifespan.

### 4.2. Proposal of a stage specific peptide cocktail strategy for myogenesis

To explain the direct cellular effects observed *in vitro* (Figure 4-5), our findings reveal that the pro-myogenic efficacy of GD is not driven by a single predominant peptide, but rather by a sophisticated, stage-specific coordination of multiple sequences. Based on the distinct temporal peaks of activity observed in our study, we propose a Multistage Synergistic Model that explains the robust efficacy of the natural GD mixture through a relay-like synergistic mechanism **(Figure 6)**.

**Figure 6.**
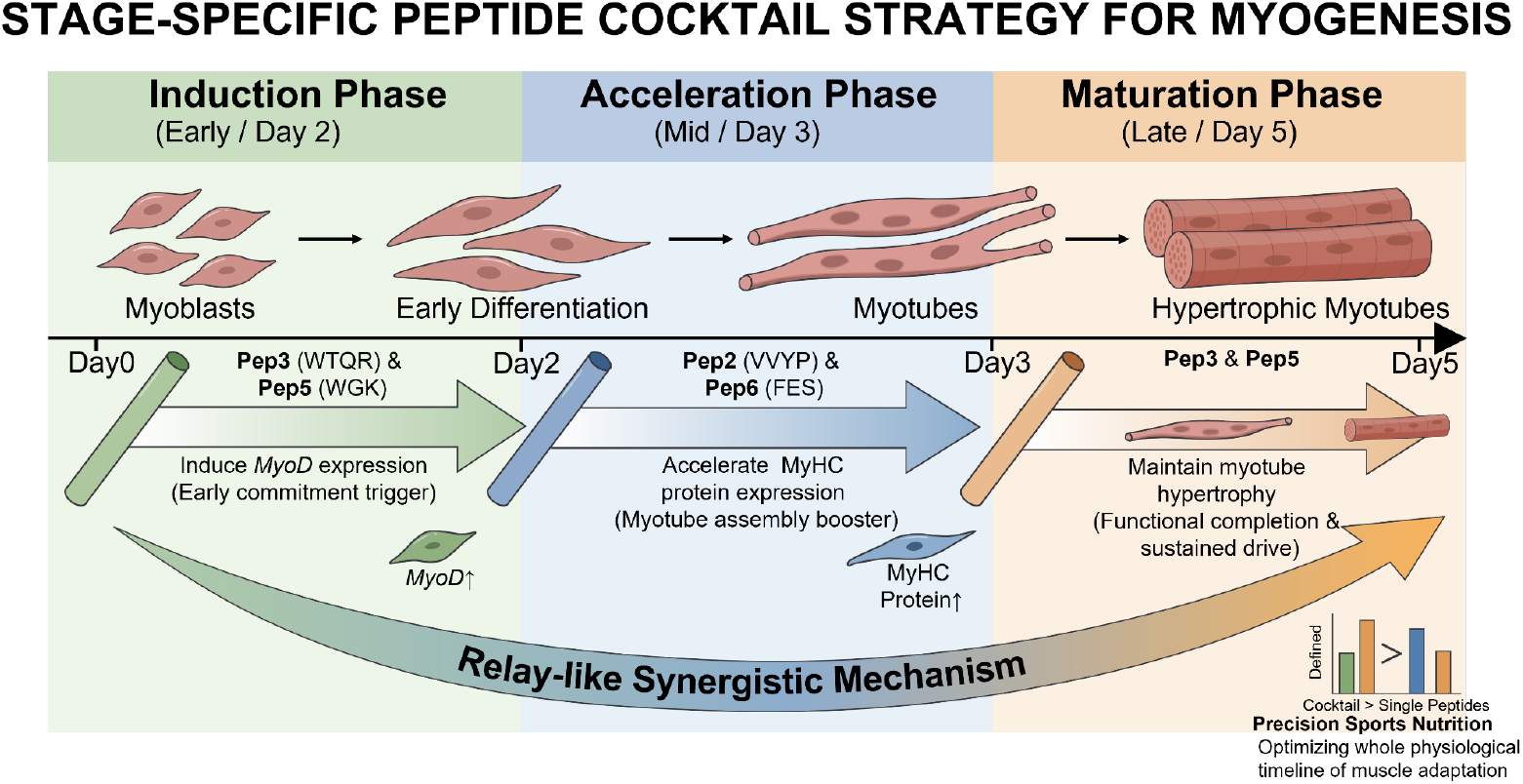
Proposed multistage synergistic model for the myogenic effects of GD peptide cocktail. The schematic illustrates the temporal, “relay-like” mechanism by which the natural GD mixture promotes skeletal muscle differentiation from myoblasts to hypertrophic myotubes. During the Induction phase (Early / day 2), Peptides 3 (WTQR) and 5 (WGK) act as early commitment triggers to potently induce *MyoD* expression. As differentiation progresses to the Acceleration phase (Mid / day 3), Peptides 2 (VVYP) and 6 (FES) function as myotube assembly boosters, accelerating MyHC protein expression. Finally, in the Maturation phase (Late / day 5), Pep 3 and 5 provide sustained drive to ensure the functional completion and maintenance of myotube hypertrophy. Natural GD Mixture achieves robust efficacy through temporal coordination, optimizing the entire physiological timeline of muscle adaptation as a targeted strategy for precision sports nutrition.

#### 4.2.1. Induction Phase (Early phase / day 2)

Interestingly, the pro-myogenic potential of GD is initiated even prior to the onset of differentiation. Our results demonstrated that GD, particularly Peptides 2 and 3, significantly stimulated myoblast proliferation (Figure S4), effectively expanding the pool of available progenitor cells. This pre-differentiation expansion, combined with the subsequent stage-specific relay mechanism, likely contributes to the robust muscle hypertrophy observed in our *in vivo* models.

Following this initial expansion, Peptides 3 and 5 function as primary triggers. Our gene expression analysis showed that these two peptides potently induce the expression of *MyoD* (Figure 5B), the master regulator of myogenic commitment. This early kick-off efficiently drives myoblasts into the differentiation program, setting a strong foundation for the subsequent stages.

#### 4.2.2. Acceleration phase (Mid phase / day 3)

Interestingly, as the process shifts toward myotube formation, Peptides 2 and 6 exhibit their peak activity at the protein level (Figure 4D). By day 3, these peptides showed significant increases in MyHC expression, suggesting they act as boosters that accelerate the translation or early assembly of the contractile apparatus, preventing any kinetic lag in the myogenic drive. Maturation phase (late phase / day 5): In the final maturation stage, the sustained myogenic drive.

#### 4.2.3. Maturation phase (Late phase / day 5)

In the final maturation stage, the sustained myogenic drive is once again anchored by Peptides 3 and 5 (Figure 4F). Their ability to maintain significantly high MyHC-positive rates through day 5 ensures the functional completion of myotube hypertrophy.

In our *in vitro* experiments, we observed an interesting dynamic regarding this late-stage maturation; while MyHC protein accumulation was significantly elevated at day 5, the mRNA levels of late-stage markers such as *Myh1* and *Myog* showed no significant differences at this same time point. This discrepancy likely reflects the temporal lag between transcription and translation; the peak of mRNA expression for these markers presumably occurred earlier, leading to the robust protein levels observed during the terminal stage of differentiation.

Overall, these results suggest that the bioactive peptides, particularly Peptides 3 and 5, may primarily accelerate the early stages of the myogenic program by modulating *MyoD* and *Myf5* expression, leading to the enhanced MyHC protein accumulation observed in later stages.

By strategically covering these temporal gaps, where early-responders (Pep3, Pep5) initiate the program, accelerators (Pep2, Pep6) maintain momentum, and sustained-drivers (Pep3, Pep5) finalize maturation, the natural GD mixture achieves a level of efficiency that any single peptide may fail to sustain. Although the direct superiority of a defined peptide cocktail over individual components remains to be empirically validated in future studies, the temporal diversity of the activity profiles observed in this study provides a highly rational model for explaining the robust efficacy of the natural GD mixture. This multistage synergistic approach represents a significant advancement toward “Precision Sports Nutrition,” where the goal is to optimize the entire physiological timeline of muscle adaptation.

### 4.3. Molecular basis for the myogenic initiation by WTQR and WGK

Within this multistage model, the early initiation of the myogenic program by Peptide 3 (WTQR) and Peptide 5 (WGK) serves as the critical first step. To understand the molecular basis of this robust induction, we focused on their structural characteristics. To date, only a few specific short-chain peptides have been recognized as direct signaling ligands. For instance, the collagen-derived dipeptides prolyl-hydroxyproline (Pro-Hyp)^[50]^ and hydroxyprolyl-glycine (Hyp-Gly)^[51]^ have been reported to promote myotube hypertrophy by modulating the Akt/mTOR/Foxo3a signaling pathway. While these collagen dipeptides primarily act to stimulate protein synthesis and suppress degradation in existing myotubes, our findings reveal that the GD-derived peptides, WTQR and WGK, exert a more profound upstream effect. By directly upregulating myogenic regulatory factors such as *MyoD* and *Myf5*, these specific GD constituents actively drive the initial commitment and differentiation of myocytes.

The unique ability of WTQR and WGK to trigger this upstream signaling is likely attributable to their specific structural characteristics, specifically the arrangement of aromatic (Trp) and basic (Arg, Lys) amino acids. This structure-activity relationship suggests that the specific amino acid sequence, rather than total nitrogen content, is the primary driver of the observed effects. The potency of these sequences may be attributed to the inclusion of tryptophan (Trp), a key regulator of muscle protein proteostasis. Recent studies indicate that serum Trp levels are positively correlated with skeletal muscle mass and that Trp is essential for maintaining muscle fiber diameter and metabolic homeostasis; its deficiency significantly impairs myoblast differentiation and muscle-specific glycolysis^[52,53]^. Furthermore, the presence of basic residues (Arg or Lys) in these sequences is critical from a nutritional physiology perspective. Recent evidence has confirmed the expression of the peptide transporter PHT1 (SLC15A4) in skeletal muscle^[51,54,55]^, which recognizes and transports short-chain peptides containing basic residues^[56–58]^. Notably, PHT1 (SLC15A4) is not merely a passive transporter but has been identified as a key scaffold protein that interacts with the Ragulator-Rag complex on the lysosomal membrane to recruit and activate mTORC1^[59,60]^. Given that the mTORC1 pathway is indispensable for the initiation of the myogenic program, the robust induction of *MyoD* and *Myf5* by WTQR and WGK observed in this study suggests that these GD-derived peptides may trigger the lysosomal signaling axis via PHT1. By potentially enhancing the translation of these early myogenic factors and triggering an autoregulatory transcriptional loop, this pathway could directly drive myocyte commitment.

### 4.4. Next-generation multifunctional peptides for unlocking the full physiological potential of athletes

The functional properties of GD and its constituent peptides elucidated in this study establish their role as hybrid bioactive molecules that transcend the scope of conventional protein supplements. Notably, the robust upstream induction of the myogenic program observed *in vitro* successfully translated into significant morphological and functional adaptations *in vivo*. Our results demonstrated that GD administration led to evident myofiber hypertrophy in mice, which directly correlated with enhanced whole-body motor performance, including increased grip strength and rotarod endurance. Furthermore, the significant improvement in stimulus-evoked burst swimming speed and acceleration in zebrafish underscores the practical impact of GD on explosive power. In our *in vivo* analysis, GD significantly upregulated the expression of fast-twitch markers (*Myh1* and *Myh2*), providing a molecular basis for these functional improvements. Given that *MyoD* and *Myf5* are critical determinants for the development of type II (fast-twitch) muscle fibers ^[8–10]^, these cross-species functional improvements from the initial trigger of *MyoD* in progenitors (Figure 5B) to the functional maturation of *Myh1/2* in tissue (Figure 3A and 3B), strongly suggest that the GD peptide relay preferentially stimulates the type II fiber-driven performance essential for high-intensity athletic activities.

Crucially, the synergistic effects are not confined to any single category of athlete. For power-based athletes pursuing hypertrophy, these peptides serve as a potent trigger for efficient muscle mass accumulation. For endurance athletes requiring rapid repair of damaged tissues^[61,62]^, GD would function as an essential recovery tool. Severe exercise-induced muscle damage (EIMD) can compromise athletic performance for several days, underscoring the urgent need for nutritional strategies—specifically bioactive peptide supplementation—to mitigate these effects. The efficacy of protein hydrolysates (WPH) in accelerating recovery is supported by robust clinical evidence, ranging from the rapid regeneration of force-generating capacity following eccentric exercise^[63]^ to the reduction of circulating creatine kinase (CK) levels and maintenance of muscle function in female populations^[64]^. By proactively driving the myogenic program—from the initial trigger of *MyoD* to the upward trend in *Myh7* expression (which is critically involved in tissue repair and oxidative function) (Figure 3D)—GD provides a universal mechanism to shorten the window required for structural tissue repair in athletes of all genders. Furthermore, for competitors in weight-class sports who must balance strict weight management with peak performance maintenance^[65]^, the simultaneous fat-reducing and muscle-sparing properties of GD provide a highly advantageous nutritional framework. Ultimately, the strategic application of this naturally derived, multistage peptide formulation represents a next-generation nutritional strategy designed to fundamentally support the physiological potential of all athletes, regardless of discipline or gender.

### 4.5. Limitations and future perspectives

While the multi-model approach employed in this study provides robust evidence for the pro-myogenic effects of GD, several limitations must be considered. First, although the identification of specific peptides that induce MyoD-mediated MyHC expression serves as a powerful strategy for muscle enhancement, human clinical trials are indispensable to confirm the clinical efficacy of the proposed peptide orchestration strategy, particularly regarding optimal dosage and timing for athletes. Furthermore, it should be noted that this study did not include a direct comparative analysis between GD and conventional intact proteins regarding gastrointestinal tolerance or net muscle protein synthesis efficiency at equivalent doses. While the low-dose efficacy of GD observed in our models is encouraging, future research is required to confirm whether GD intake effectively mitigates digestive distress while maintaining peak performance compared to standard high-protein diets.

## 5. Conclusion

In conclusion, this study establishes GD and its constituent peptides as potent signaling modulators that transcend the traditional role of protein as a mere building block. By demonstrating that a specific “peptide relay” directly activates the myogenic program and modulates the adipo-muscular axis, GD offers a targeted strategy for “Precision Sports Nutrition” to optimize performance and a novel approach to combat sarcopenic obesity. These findings provide a robust foundation for unlocking the full physiological potential of the human body to support lifelong functional vitality.

## Abbreviations

Akt: Protein Kinase B (PKB)
AMPK: AMP-activated Protein Kinase
BDNF: Brain derived neurotrophic factor
C2C12: C2C12 myoblasts
EIMD: Exercise-induced muscle damage
FIHC: Fluorescent immunohistochemistry
Foxo3a: Forkhead box O3
GD: Globin digest
GLP-1: Glucagon-Like Peptide-1
HE: Hematoxylin and Eosin
MyHC: Myosin heavy chain
Mtor: Mammalian target of rapamycin
Myf5: Myogenic factor 5
*Myh1* / *Myh2* / *Myh3* / *Myh7*: Myosin heavy chain 1/2/3/7
*MyoD*: Myogenic differentiation 1
*Myog*: Myogenin
MRFs: Myogenic regulatory factors
mTORC1: mTOR Complex 1
mpf: months post fertilization
Pro-Hyp: Prolyl-hydroxyproline
Hyp-Gly: Hydroxyprolyl-glycine
PRKAA2: Protein kinase AMP-activated catalytic subunit alpha 2
SLC15A4: Solute Carrier Family 15 Member 4
UCP1: Uncoupling protein-1
UCP2/3: Uncoupling protein 2 / 3
PepT1: Peptide transporter 1
PHT1: Peptide/histidine transporter 1

## Author Contributions

K.F., N.N., and Y.S. designed the study. M.N., L.Z., and Y.S. performed the experiments. K.F. and K.I. provided essential resources. M.N. and Y.S. wrote the manuscript. L.Z., K.F., and Y.S. reviewed and edited the manuscript.

