## Supplemental Table and Figures for "Globin digest and its constituent peptides promote skeletal muscle hypertrophy and enhance physical performance"

### Supplementary Materials

**Table S1. Primer sequences used for qRT-PCR analysis.**

| Gene | Forward (5'-3') | Reverse (5'-3') | Reference |
|---|---|---|---|
| *Hprt* | ATGGACTGATTATGGACAGGACTG | TCCAGCAGGTCAGCAAAGAAC | [33] |
| *Gapdh* | GTTGTCTCCTGCGACTTCA | GGTGGTCCAGGGTTTCTTA | [33] |
| *Myh1* | GAGGGACAGTTCATCGATAGCAA | GGGCCAACTTGTCATCTCTCAT | [34] |
| *Myh2* | AGGCGGCTGAGGAGCACGTA | GCGGCACAAGCAGCGTTGG | [34] |
| *Myh3* | TCCGACAACGCCTACCAGTT | CCCGGATTCTCCGGTGAT | [34] |
| *Myh7* | ATGAGCTGGAGGCTGAGCA | TGCAGCCGCAGTAGGTTCTT | [34] |
| *Mtor* | AGAAGGGTCTCCAAGGACGACT | GCAGGACACAAAGGCAGCATTG | [35] |
| *Myog* | CAGCCCAGCGAGGGAATTTA | AGAAGCTCCTGAGTTTGCCC | [36] |
| *MyoD* | AGCACTACAGTGGCGACTCA | GGCCGCTGTAATCCATCA | [37] |
| *Myf5* | CTGCTCTGAGCCCACCAG | GACAGGGCTGTTACATTCAGG | [38] |

**Figure S1**

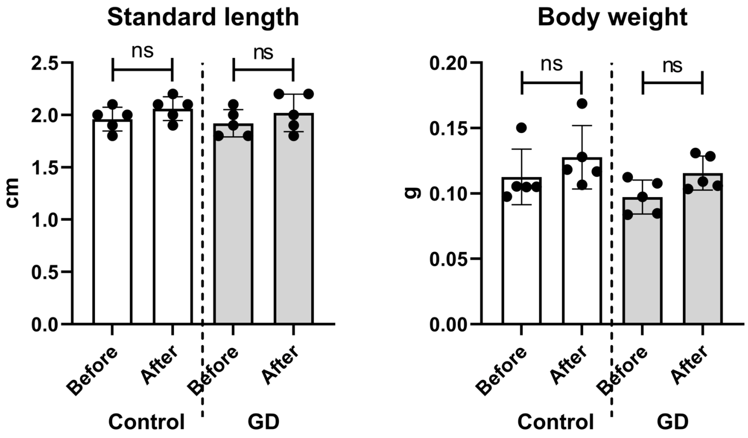


**Figure S1. Effects of short-term GD administration on the physical growth of adult zebrafish.** Changes in standard length (left panel) and body weight (right panel) of zebrafish before and after the 1-week administration period in the control and GD-treated groups. Data are presented as mean ± standard deviation (SD). ns: not significant. Control: n = 5; GD: n = 5.

**Figure S2**

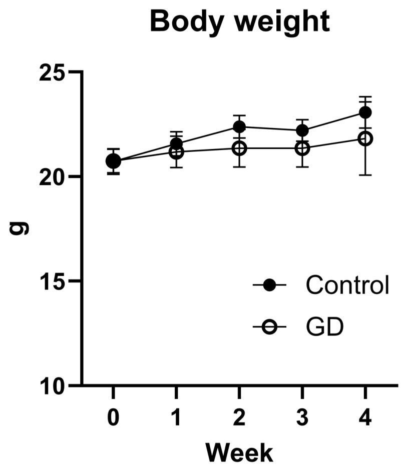


**Figure S2. Effects of GD administration on body weight in mice**. Body weight changes were monitored weekly during the 4-week administration period. Data are presented as mean ± SD. No statistically significant differences were observed between the control and GD-treated groups at any time point (n = 6 per group).

**Figure S3**

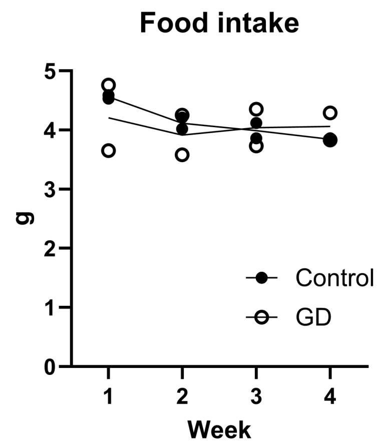


**Figure S3. Effects of GD administration on average food intake in mice.** Average daily food intake per mouse is shown over the 4-week administration period. Food consumption was measured per cage (3 or 4 mice per cage), and the weekly average was calculated. Data are presented as mean values with individual data points for each cage. No apparent differences were observed between the control and GD-treated groups (n = 2 cages per group).

**Figure S4**

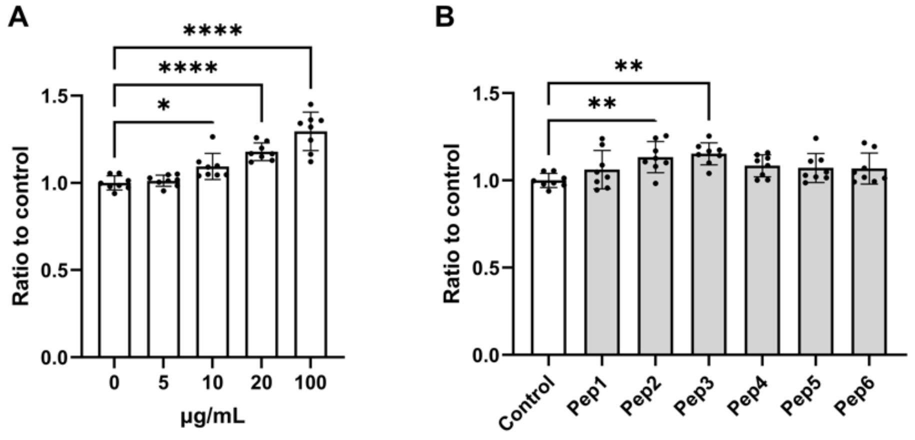


**Figure S4. Effects of GD and its constituent peptides on myoblast proliferation.** Undifferentiated C2C12 myoblasts were cultured in growth medium and treated with the indicated compounds for 48 h. Cell proliferation was evaluated using a CellTiter-Glo Luminescent Cell Viability Assay (Promega, Madison, WI, USA). (**A**) Dose-dependent effects of GD on cell proliferation. Cells were treated with GD at concentrations of 5, 10, 20, and 100 µg/mL. (**B**) Effects of the six individual constituent peptides on cell proliferation. Each synthetic peptide was administered at a concentration of 1 µg/mL. Data are expressed as a ratio to the control group (0 µg/mL) and presented as mean ± SD with individual data points (n = 8 per group). *$p < 0.05$, **$p < 0.01$, ****$p < 0.0001$ vs. control.
